# Network simulations of interneuron circuits in the honeybee primary auditory center

**DOI:** 10.1101/159533

**Authors:** Ajayrama Kumaraswamy, Aynur Maksutov, Kazuki Kai, Hiroyuki Ai, Hidetoshi Ikeno, Thomas Wachtler

**Affiliations:** Department of Biology II, Ludwig-Maximilians-Universität München, 82152 Planegg-Martinsried, Germany; Department of Biomedical Physics, Moscow Institute of Physics and Technology, Dolgoprudny, Russia; Department of Earth System Science, Fukuoka University, 814-0180 Fukuoka, Japan; School of Human Science and Environment, University of Hyogo, 670-0092 Himeji, Japan

## Abstract

Processing of airborne vibration signals in the auditory system is essential for honeybee communication through the waggle dance language. Properties of neurons in the honeybee primary auditory center suggest a circuitry of excitatory and inhibitory neurons encoding these communication signals. To test this assumption, we simulated this network and analyzed the predicted responses for different types of inputs. In particular, we investigated the effect of specific inhibitory connections in the network. The results indicate that the experimentally observed responses of certain interneuron types are compatible with an inhibitory network of vibration processing in the primary auditory center of the honeybee.

## INTRODUCTION

### Waggle dance and the honeybee primary auditory center

One of many fascinating behaviors of the honeybee is the “Waggle Dance”, which is used by returning forager honeybees to advertise the location of beneficial resources like food, water and pollen among hive mates (Von Frisch 1967). The waggle dance consists of the “waggle phase” during which the honeybee walks in a specific direction wagging its body and flapping its wings, and a “return phase” during which it returns along a curved path to the start of the waggle phase. The waggle phase represents the flight path of the bee from the hive to resource, encoding the distance and direction of the resource in its duration and orientation, respectively. During the waggle phase, body and wing movements of the dancer honeybee produce pulses of air vibrations of specific frequency (≈265 Hz), duration (≈16 ms) and inter-pulse-interval (≈33 ms) (Wenner 1962). These “sounds” are important for successful recruitment of foragers (Barth et al. 2005; Michelsen 2003) and are sensed by follower bees using the Johnston’s Organ (JO) in their antennae. Sensory neurons in the JO transduce air vibrations into neural signals (Tsujiuchi et al. 2007) and convey them to the honeybee brain, specifically to the Dorsal Lobe (DL), the dorsal Sub Esophageal ganglion (dSEG) and the medial Posterior Protocerebral Lobe (mPPL) (Ai 25 et al. 2009), the regions forming the primary auditory center (PAC) of the honeybee brain.

### Experimental results about PAC neurons

Experimental studies have identified several vibration-sensitive neurons in the honeybee PAC and characterized their morphological projections and physiological responses along with those of the JO sensory afferents (Ai et al. 2009; Ai et al. 2016; Ai et al. in prep.). The JO sensory afferents that project into the DL responded to sinusoidal vibration stimuli by producing spikes with high probability at a fixed phase of the input (unpublished data). DL-Int-1, a local inhibitory interneuron that arborizes in the DL, dSEG and mPPL, showed close proximity to JO sensory afferents (Ai and Hagio 2013) and responded to one-second-long continuous sinusoidal vibration with on-phasic excitation and tonic inhibition followed by post-inhibitory rebound (Ai et al. 2009). DL-Int-2, an excitatory output neuron that arborizes in the DL, dSEG, central PPL and the lateral protocerebrum (LP), also showed close proximity to JO sensory afferents (Ai and Hagio 2013). It responded to one-second-long continuous sinusoidal vibration with on-phasic and tonic excitation. For stimulation with trains of vibration pulses with different pulse duration and inter-pulse-interval (IPI) values (Ai et al. 2016; Ai et al. in prep.), DL-Int-1 showed a reduced spiking rate caused by strong inhibition for pulse trains of IPI shorter than 33 ms, which gradually increased for stimulus trains of higher IPIs. In contrast, DL-Int-2 showed higher spiking rates for pulse trains of IPI up to 33 ms, which reduced for pulse trains of higher IPI.

### Putative Network model

The response properties, together with projection patterns and immunohistochemistry, suggested a disinhibitory network involving DL-Int-1 and DL-Int-2 (Fig 1A; Ai et al. in prep.). In this model, sensory afferents from the JO are assumed to form excitatory synapses onto both DL-Int-1 and DL-Int-2. The delayed tonic inhibition shown by DL-Int-1 is assumed to be the result of a local inhibitory neuron that is driven by JO sensory afferents. Since DL-Int-1 is GABAergic and has boutons in the DL and dSEG where DL-Int-2 arborizes (Ai et al. 2009), an inhibitory synapse is assumed from DL-Int-1 to DL-Int-2.

**Figure 1.**
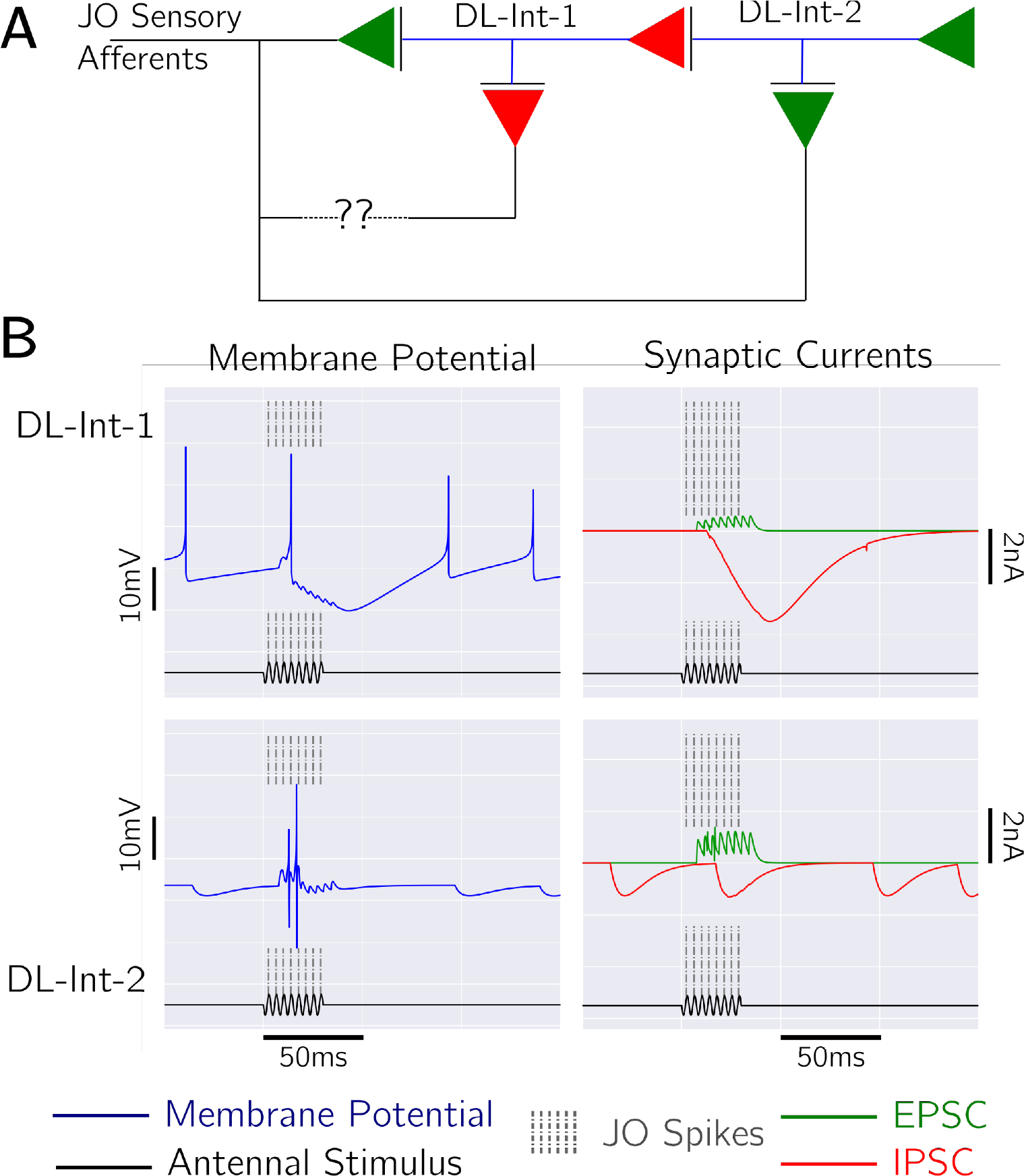
**A**, The network model used for the simulation. DL-Int-1 and DL-Int-2 were modeled as point neurons using the Adaptive Exponential Model and the synaptic conductances were modeled as difference of exponentials. **B**, Example simulation traces for a continuous sinusoidal stimulus of length 30 ms and frequency 265 Hz. The membrane potential traces for DL-Int-1 and DL-Int-2 are shown in the left column with their excitatory and inhibitory synaptic currents in the right column.

To test these assumptions and investigate the role of inhibition and disinhibition in vibration processing in the honeybee, we used phenomenological neuron models of DL-Int-1 and DL-Int-2 and simulated different interneuron circuits with variants of inhibitory connections.

## Methods

### Choice of neuron and synapse models

Since very little is known about the membrane properties of DL-Int-1 and DL-Int-2, point neuron models were chosen instead of more detailed models. The AdExp model is well understood (Touboul and Brette 2008) and can replicate a wide variety of neural responses with few parameters (Rossant et al. 2011; Kremer et al. 2011; Vogels et al. 2011). The double-exponential synaptic conductance model (Carnevale and Hines 2006) is a good approximation for experimentally observed synaptic traces (e.g.: Häusser and Roth 1997; Zsiros and Hestrin 2005). It provides parameters to independently control the rise and fall time constants and synaptic strength, which was specifically useful to implement the single-neuron properties of DL-Int-1 and DL-Int-2.

### Model implementation

JO sensory neurons were assumed to spike at the positive peak of the input sinusoidal vibration stimulus. DL-Int-1 and DL-Int-2 were modeled as point neurons with their membrane potentials simulated using the Adaptive Exponential Integrate-and-fire (AdExp) model (Naud et al. 2008) (Fig 2). These membrane potential calculations included synaptic input currents, which which were calculated from their conductances. Synaptic conductances were simulated using difference of exponentials functions, based on the Exp2Syn model of the simulator NEURON (Carnevale and Hines 2006; `RRID:SCR_005393`) (Fig 3).

**Figure 2.**
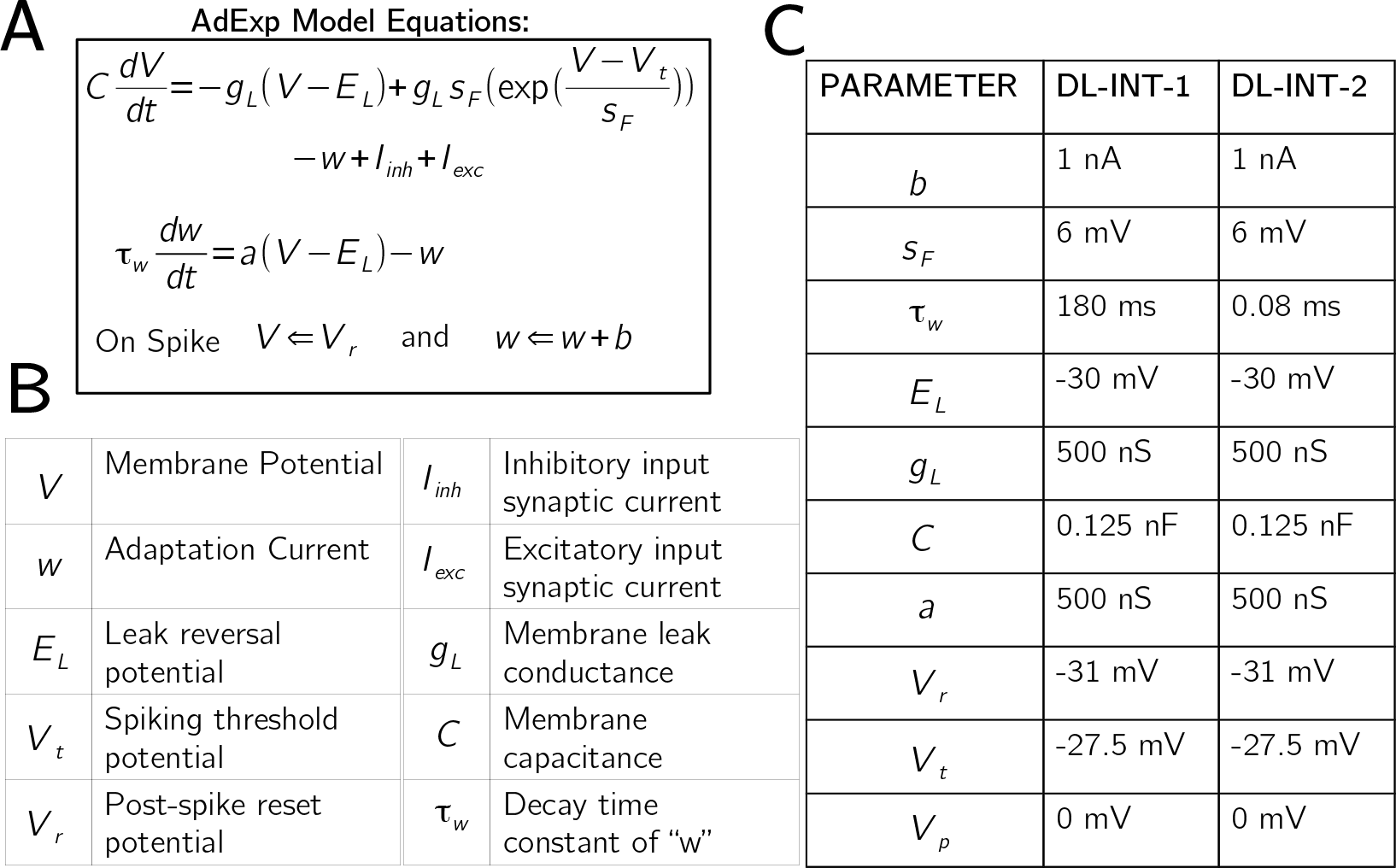
**A**, Equations of the Adaptive Exponential integrate-and-fire (AdExp) point
neuron model used for DL-Int-1 and DL-Int-2. **B**, Description of the parameters of the equations in A. **C**, The set of parameters that were used in the simulations.

**Figure 3.**
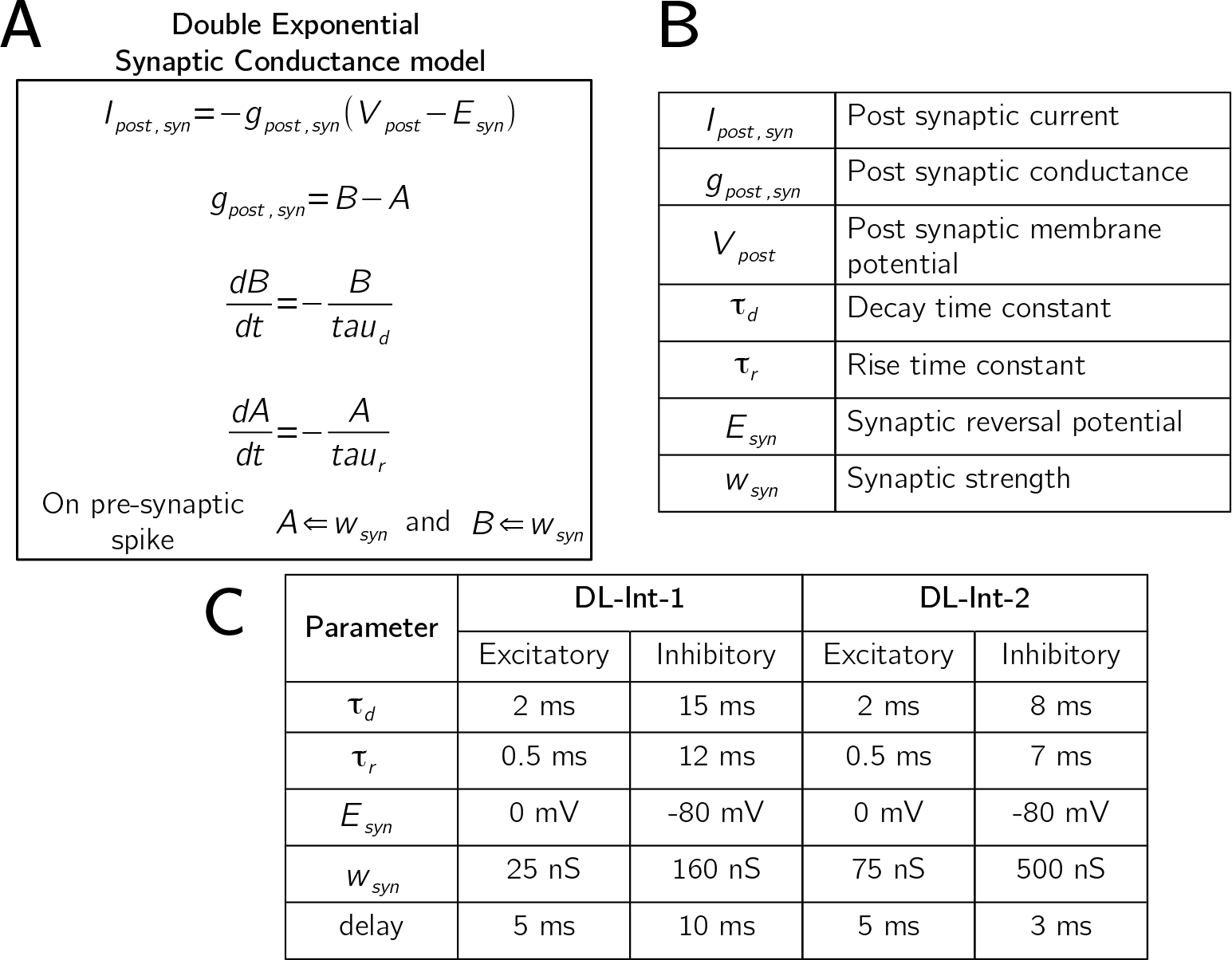
**A**, Equations of the difference of exponentials synaptic conductance model used
for the excitatory and inhibitory synapses. **B**, Description of the model parameters. **C**, The set of parameters that were used for simulations.

### Simulation setup and stimuli used

The network model described above was implemented using the simulator Brian version 2.0.1 (Stimberg et al. 2014;`RRID:SCR_002998`) in Python. An integration step size of 0.1 ms was used and all simulation runs had a “settling time” of 600 ms, after which stimuli were applied. Similar to experimental studies (Ai et al. 2009; Ai et al. 2016; Ai et al. in prep.), continuous sinusoidal stimuli with stimulus frequency 265 Hz and different stimulus durations were used, along with trains of sinusoidal pulses of stimulus frequency 265 Hz and different combinations of pulse durations and IPI values. The pulse parameters were chosen around the waggle dance parameters of 16ms pulse duration and 33 ms IPI.

The code used to simulate this network is available on Github at https://github.com/wachtlerlab/HB-PAC_disinhibitory_network.

### Tuning model parameters

Parameters of the DL-Int-1 model and its inputs synapses were chosen to qualitatively reproduce the response characteristics of DL-Int-1. In particular, membrane and adaptation parameters of DL-Int-1 were tuned to produce non-zero spontaneous firing rate. The temporal dynamics of excitatory and inhibitory input synapses were adjusted by controlling their rise and fall time constants and their delays to qualitatively reproduce the temporal pattern of DL-Int-1's response to continuous sinusoidal pulses, i.e., its on-phasic excitation, tonic inhibition and rebound. Further, the strengths of input synapses along with the parameters that affect the neuron’s excitability, viz., *g_L_* and *a*, were adjusted to qualitatively reproduce the response patterns of DL-Int-1 to trains of vibration pulses while retaining its tuned response to continuous pulses. Parameters of DL-Int-2 were assumed to be same as DL-Int-1 except for one adaptation parameter, *τ_w_*, which was adjusted to account for the zero spontaneous firing rate and on-phasic and tonic excitation properties of this neuron type. The excitatory input synapse of DL-Int-2 had the same temporal dynamics as those of DL-Int-1, while its inhibitory input synapse was modified to have faster dynamics. The strengths of these synapses were tuned to qualitatively replicate DL-Int-2’s response properties to trains of vibration pulses. All tuning parameters are listed along with their descriptions in Figs. 2 (neuron parameters) and 3 (synapse parameters).

## Results

### Qualitative replication of experimental results

Results for simulations of the neurons with a continuous sinusoidal vibration stimulus of 1 s are shown in Fig 4. The simulated membrane potential traces of DL-Int-1 showed non-zero spontaneous activity, on-phasic excitation, tonic inhibition and rebound, qualitatively very similar to experimental traces (see Figs 6 and 7 of Ai et al. 2009). The simulated membrane potential changes of DL-Int-2 showed zero spontaneous activity, on-phasic and tonic inhibition.

**Figure 4.**
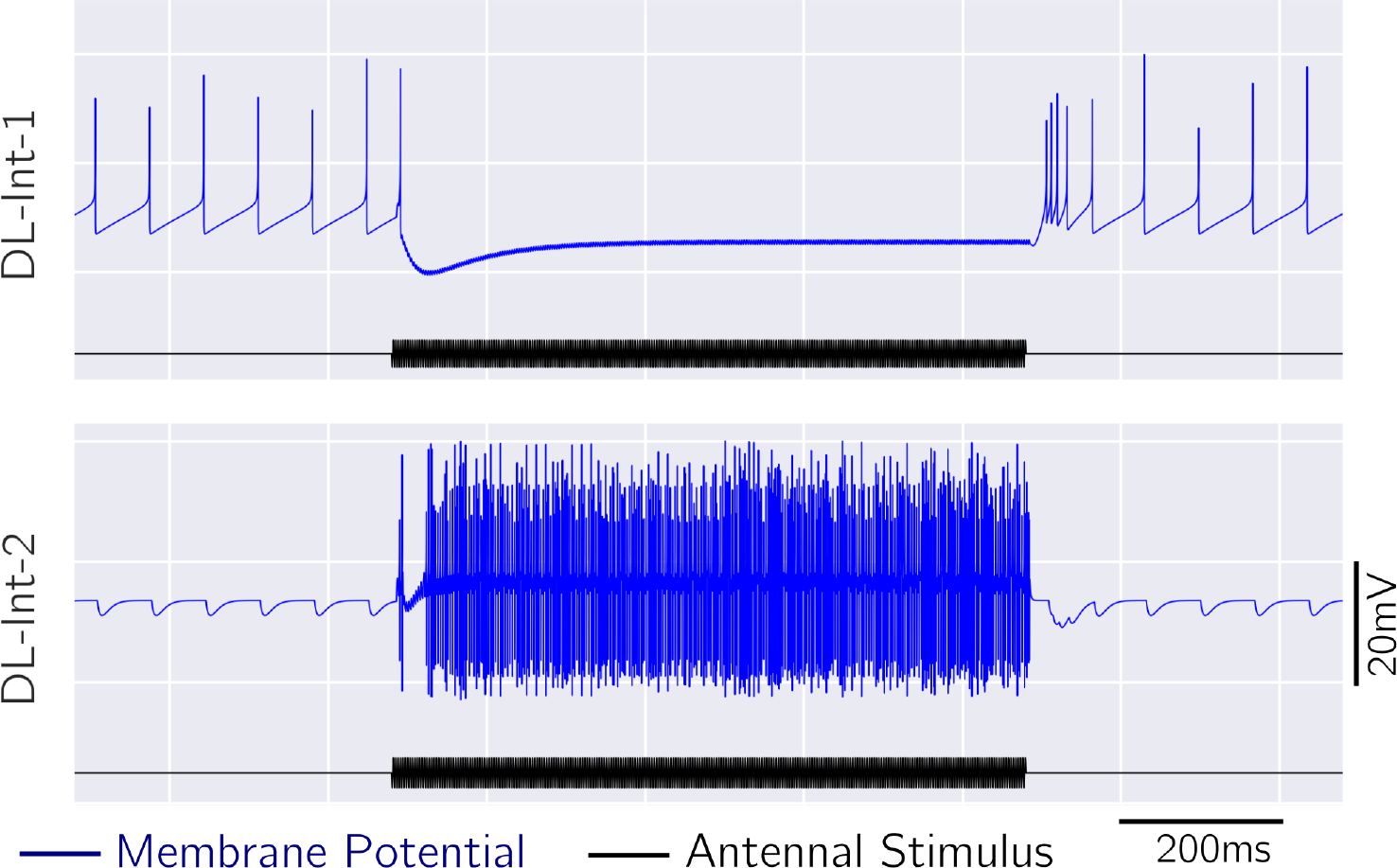
Responses of DL-Int-1 and DL-Int-2 to continuous sinusoidal vibration stimuli of frequency 265Hz. DL-Int-1 showed non-zero spontaneous activity, on-phasic excitation and tonic inhibition followed by post inhibitory rebound. DL-Int-2 showed zero spontanous activity, on-phasic and tonic excitation.

Membrane potential traces of DL-Int-1 for different trains of sinusoidal pulses are summarized in Fig. 5. Simulations showed low spiking rates for pulse trains of IPI shorter than 33 ms and increasingly higher spike rates for pulse trains with IPIs above 33 ms.

**Figure 5.**
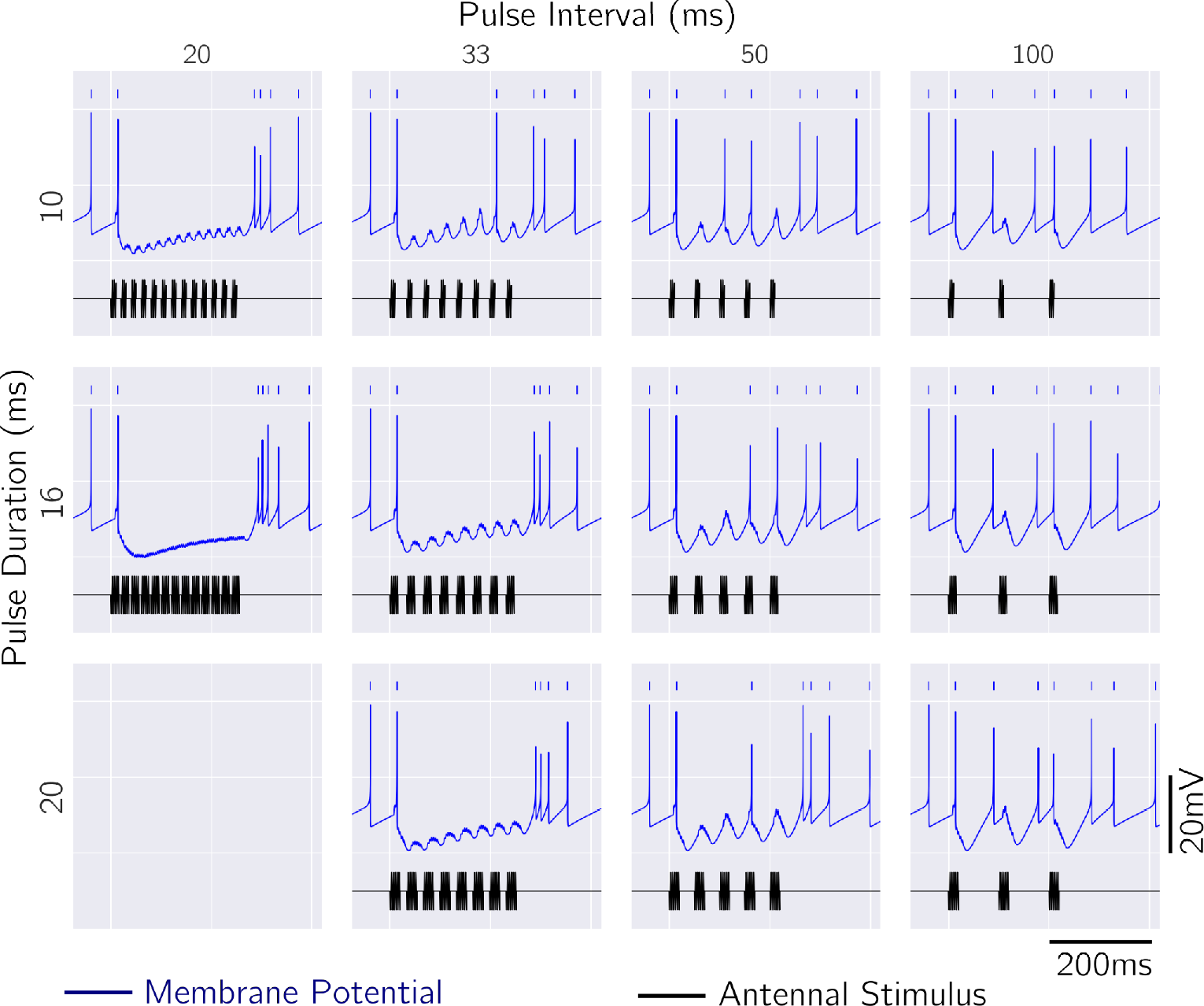
Simulated responses of DL-Int-1 to trains of vibration pulses of different pulse
durations and pulse intervals. Spikes corresponding to the blue membrane potential traces are shown using short blue vertical lines. The model showed very low spiking rates for lower pulse intervals and higher spiking rates for higher pulse intervals. Increasingly stronger sub-threshold oscillations were observed for increasing pulse interval values.

### Significance of Inhibitory synapse from DL-Int-1 to DL-Int-2

A summary of simulated membrane potential traces of DL-Int-2 in response to pulse trains with different pulse parameters is shown in Fig. 6 (blue). DL-Int-2 responded to pulse trains with IPIs shorter than 33 ms with high spike rates, which reduced for trains with longer IPIs. The significance of the inhibitory connection from DL-Int-1 to DL-Int-2 was tested by silencing this inhibitory input in the model. With the inhibitory input from DL-Int-1 absent, DL-Int-2 showed high firing rates for all pulse parameters (Fig6, red). Since DL-Int-1 had low firing rate for pulse trains with short intervals, it did not affect the response of DL-Int-2 for these input parameters. However, DL-Int-1 showed higher spiking rate for pulse trains of IPI greater than 33 ms and hence could affect the response of DL-Int-2.

**Figure 6.**
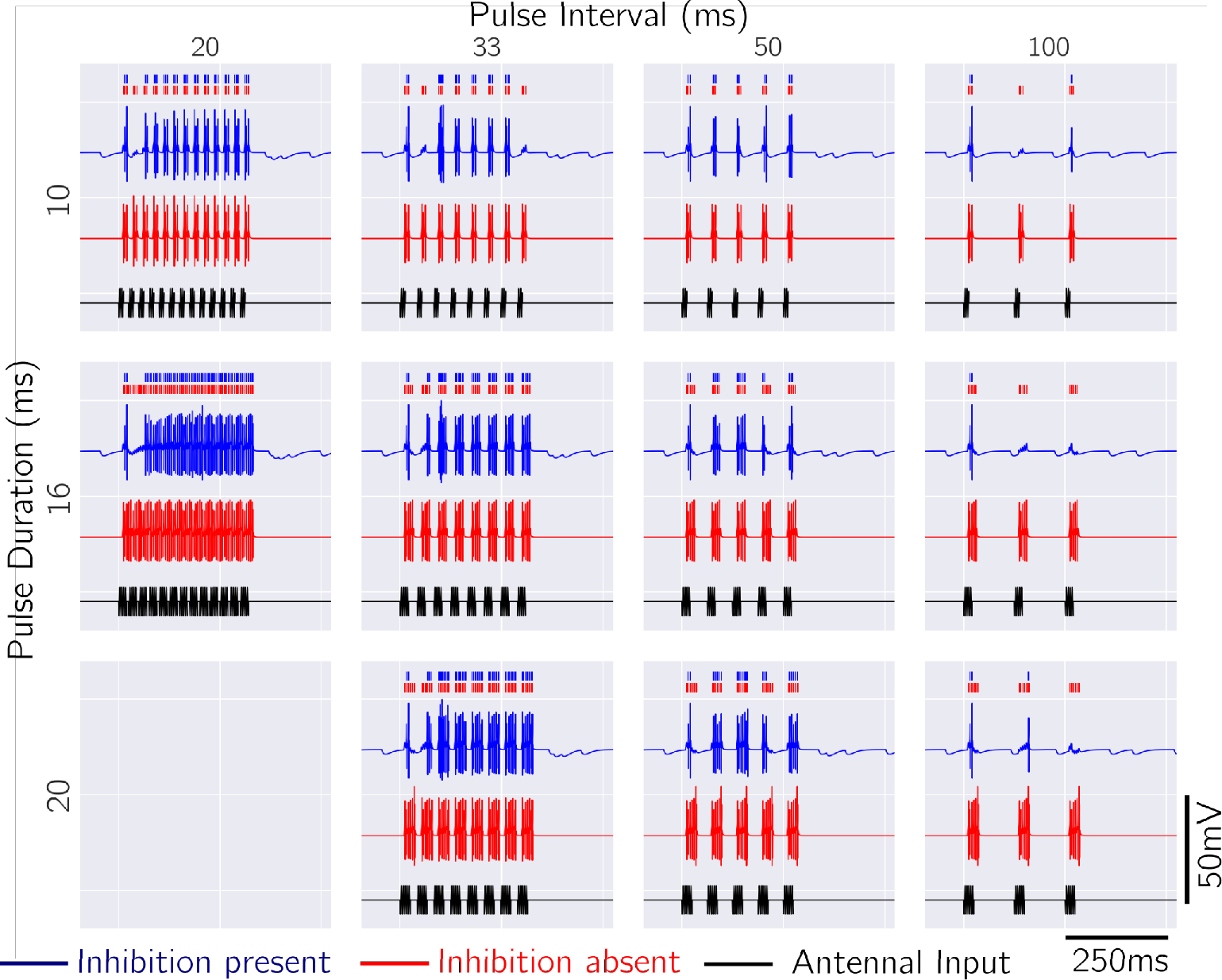
Simulated responses of DL-Int-2 to trains of vibration pulses of different pulse
intervals and durations for the cases where the inhibition from DL-Int-1 is present (blue) and absent (red). Spikes corresponding to the membrane potential traces are shown using short vertical lines of corresponding colors. With the DL-Int-1 inhibition present (blue), the model showed high firing rate for short pulse intervals which gradually reduced for higher pulse intervals, consistent with experimental observations. Such a reduction in firing rate for higher pulse intervals is not seen for the case without DL-Int-2 inhibition (red).

## Discussion

In this study, we simulated a hypothesized network of interneurons in the honeybee primary auditory center to test its consistency with experimentally observed responses. Simulations qualitatively reproduced the membrane potential traces of the interneurons to continuous vibration stimuli as well as trains of vibration pulses, suggesting that the hypothesized network model could underlie the experimental observations.

Since the simulation results of this study are only qualitative, and in particular the parameters of the model neurons were not directly measured experimentally, the results confirm the consistency, but do not imply the necessity, of the assumed circuitry. For example, the same behavior might potentially be observed in a network with different circuitry, possibly requiring different neuron properties. However, building and testing such models would require more experimental data.

This simulation study is a first step towards understanding the neural circuitry that underlies auditory processing in the honeybee. Although the simulated model provides instructive insights like the level of required excitability of the interneurons and the interplay between excitatory and inhibitory synaptic currents for producing the observed experimental traces, they do no explain all the observed features of experimental responses to all the stimulus patterns used. More extensive testing and comparison of more complex models would be required, which however would require more information about synaptic connectivity and individual membrane properties of the interneurons of the honeybee primary auditory center.

## Conclusion

We have simulated a network model of identified neurons in the primary auditory center of the honeybee brain, to investigate the potential role of inhibitory connections and in particular to test the assumption of a disihibitory network of auditory processing in the honeybee. The results show that such a network model is compatible with the experimental data. The principles underlying the network model could help to better understand auditory processing in the honeybee brain.

## Acknowledgments

Supported by the Japanese-German Collaborative Program on Computational Neuroscience (JST and BMBF grant 01GQ1611), the Bernstein Center for Computational Neuroscience Munich (BMBF grant 01GQ1004A), and the Amgen Scholars Programme 2016.

